# Molecular mechanism underlying venation patterning in butterflies

**DOI:** 10.1101/2020.01.02.892760

**Authors:** Tirtha Das Banerjee, Antónia Monteiro

## Abstract

The mechanism of wing vein differentiation in *Drosophila* is a classic text-book example of pattern formation using a system of positional-information, yet very little is known about how this mechanism differs in species with a different number of veins and how insect venation patterns evolved. Here, we examine the expression patterns of genes previously implicated in vein differentiation in *Drosophila* in two butterfly species with more complex venation, the African squinting bush brown *Bicyclus anynana* and the Asian cabbage white, *Pieris canidia*. We also test the function of one of these genes, *spalt (sal)*, with CRISPR-Cas9 in *B. anynana*. We identify both conserved as well as new domains of *decapentaplegic (dpp), engrailed (en)*, *invected (inv)* and *sal* gene expression in *B. anynana*, and propose how the simplified venation in *Drosophila* might have evolved via loss of *dpp* and *sal* gene expression domains, silencing of vein inducing programs at Sal-expression boundaries, and changes in gene expression of vein maintenance genes.

**Summary statement:** The paper describes new domains of venation patterning genes in butterflies and proposes how simplified venation in other insect lineages might have evolved.

## Introduction

Current venation patterns in several insect groups appear to be simplified versions of more complex ancestral patterns. The fossil record indicates that ancestral holometabolous insects, such as *Westphalomerope maryvonneae*, had highly complex vein arrangements which evolved into simpler venation with enhanced efficiency to sustain powered flight in modern representatives of Diptera and Lepidoptera (*1*). To identify these simplifications, Comstock and Needham in the 1900s developed a system of vein homologies across insects. The system nomenclature recognizes six veins protruding from the base of the wings called Costa (C), Sub-costa (Sc), Radius (R), Media (M), Cubitus (Cu) and Anal (A)(*2*). These veins can later branch into smaller veins, and cross-veins add additional complexity to venation patterns. Every longitudinal insect wing vein, however, can be identified using this nomenclature. Vein simplifications over the course of evolution have happened either via fusion of veins or disappearance of particular veins (*3*–*6*), but the molecular mechanisms behind these simplifications remain unclear.

Molecular mechanisms of vein pattern formation have been primarily investigated in the model vinegar fly *Drosophila melanogaster*, where a classic system of positional information takes place. Here, the wing is initially sub-divided into two domains of gene expression, and anterior and posterior compartment, where *engrailed (en)* and *invected (inv)* expression are restricted to the posterior compartment (*7*, *8*). A central linear morphogen source of the protein Decapentaplegic (Dpp) is established at the posterior border of the anterior compartment, and genes like *spalt* (*sal*) respond to the continuous morphogen gradient in a threshold-like manner, creating sharp boundaries of gene expression that provide precise positioning for the longitudinal veins (*9*, *10*). Veins differentiate along these boundaries, along a parallel axis to the Dpp morphogen source (*11*, *12*). Vein cell identity is later determined by the expression of genes such as *rhomboid (rho)*, downstream of *sal* (*10*, *13*). Conversely, intervein cells will later express *blistered* (*bls*) which suppresses vein development (*14*, *15*). The final vein positions are determined by the cross-regulatory interaction of *rho* and *bls*.

The mechanisms underlying venation patterning in other insect lineages have remained poorly understood, and so far, gene expression patterns and functions for the few genes examined seem to be similar to those in *Drosophila* (*16*–*18*).

Venation patterning in butterflies (order: Lepidoptera) has been examined in a few mutants in connection with alterations of color pattern development (*19*, *20*) and more directly via the expression pattern of a few genes during larval development. Two of the species in which a few of the venation patterning genes have been studied in some detail are the African squinting bush brown butterfly, *Bicyclus anynana* and common buckeye butterfly, *Junonia coenia*. In both species, En and/or Inv were localized in the posterior compartment using an antibody that recognizes the epitope common to both transcription factors (*21*, *22*). The transcript of *Inv* in *Junonia*, however, appears to be absent from the most posterior part of the wings (*23*), whereas the transcript of *hedgehog (hh*), a gene is up-regulated by En/Inv (*24*) has been shown to be uniformly present in the posterior compartment in both species (*21*, *25*). There is little knowledge of the expression domains of the other genes, including the main long-range morphogen *dpp* and its downstream targets (e.g., *sal*). A recent report proposed the presence of a second *dpp*-like organizer at the far posterior compartment in butterflies (*26*). This report, however, showed no direct gene expression or functional evidence and has been debated by other researchers (*27*). Currently there is also no functional evidence of altered venation for knock-out phenotypes for any of these genes in Lepidoptera.

Here we explore the expression of an orthologous set of genes to those that are involved in setting up the veins in *Drosophila* in two butterfly species: *Bicyclus anynana* and *Pieris canidia*. Then we perform CRISPR-Cas9 to test the function of one of these genes in venation patterning in *Bicyclus*.

## Results

### Expression of *engrailed* and *invected* transcripts and proteins

We first examined the expression pattern of En and Inv at both the transcript and protein levels in *Bicyclus anynana* and in *Pieris canidia* fifth instar larval wings. We used an antibody that recognizes the epitope of both proteins and confirmed that En and/or Inv expression is found throughout the posterior compartment in both forewings and hindwings in *Bicyclus* (*21*, *28*), and also in *Pieris*, however, a sharp drop in expression levels is observed posterior to the A2 vein in both species (Fig. 1, A-D). We hypothesized that the low En/Inv posterior expression could either be due to lower levels of transcription or translation of En and/or Inv, or due to the absence of either of the two transcripts in the area posterior to the A2 vein. To test these hypotheses, we performed *in-situ* hybridization using probes specific to the transcripts of *en* and *inv* in *Bicyclus* (see Supp file for sequences). *en* was expressed homogeneously throughout the entire posterior compartment on both the forewing and the hindwing, but *inv* was restricted to the anterior 70-75% of the posterior compartment (Fig. 1, E-H). Hence, the low levels of En/Inv protein expression appear to be due to the absence of *inv* transcripts in the most posterior region of the posterior compartment.

**Fig. 1.**
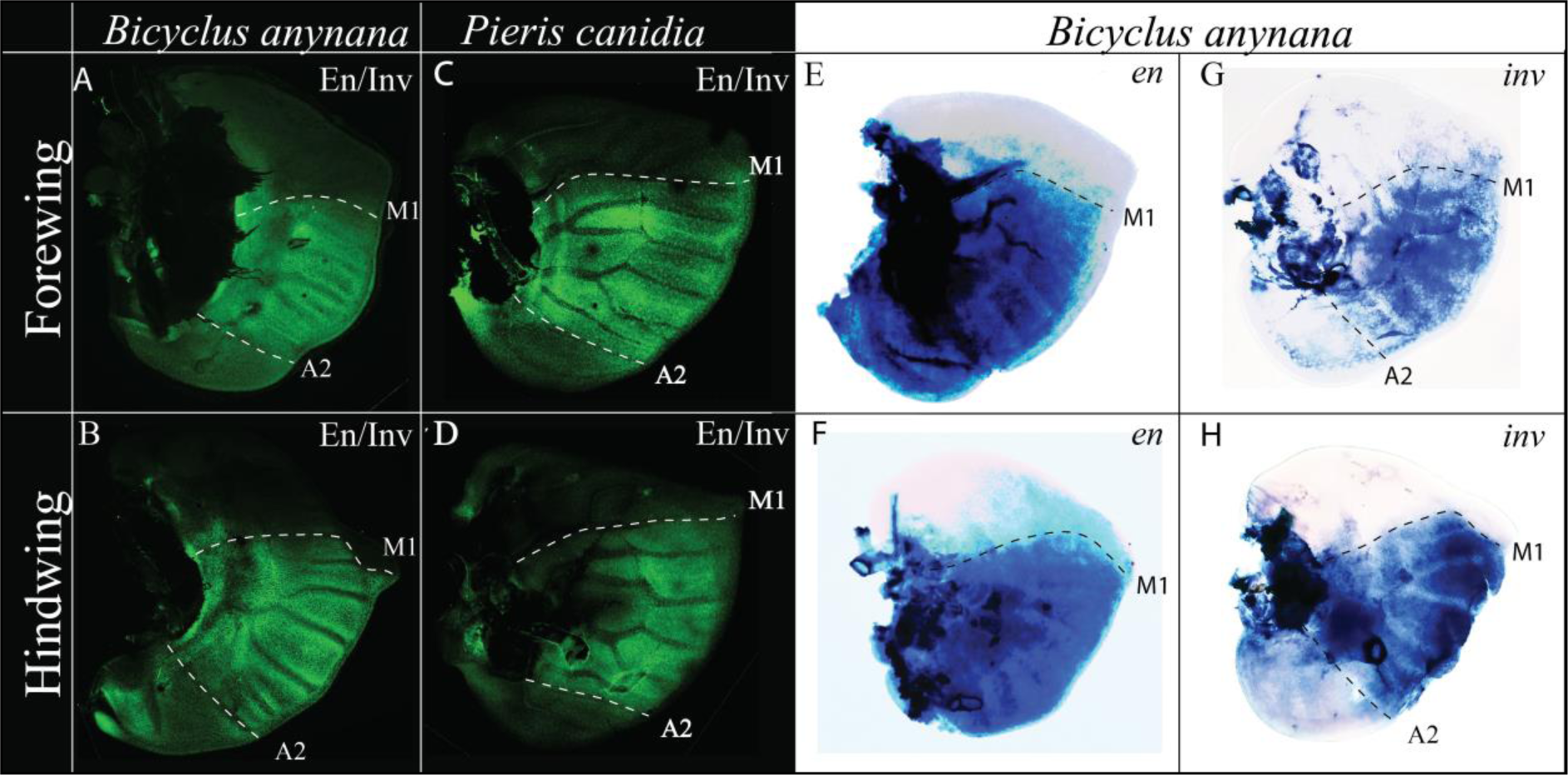
Expression of Engrailed and Invected proteins in *Bicyclus anynana* and *Pieris canidia* and expression of mRNA transcripts in *Bicyclus anynana*. **(A)** Expression of En/Inv proteins in the forewing, and **(B)** hindwing is strong between the M1 and A2 veins, and weaker posterior to the A2 vein. **(C)** Expression of En/Inv in the forewing, and **(D)** hindwing in *Pieris* is strong between the M1 and A2 veins, and weaker posterior to the A2 vein. **(E)** Expression of *en* mRNA transcripts in the forewing, and **(F)** hindwing in *Bicyclus* is almost homogeneous across the posterior compartment. **(G)** Expression of *inv* in the forewing, and **(H)** hindwing in *Bicyclus* is detected around 70% of the posterior compartment from the M1 vein till the A2 vein.

### Expression of *dpp*

We explored the presence of transcripts for the BMP signaling ligand Decapentaplegic (Dpp) with the help of *in-situ* hybridization using a probe specific to its transcript (see Supp file for sequence). A strong band of *dpp* was observed along the A-P boundary (i.e., along the M1 vein). However, another strong domain was observed in the lower posterior compartment around the A3 vein (Fig. 2A; Fig. S3, A-C).

**Fig. 2.**
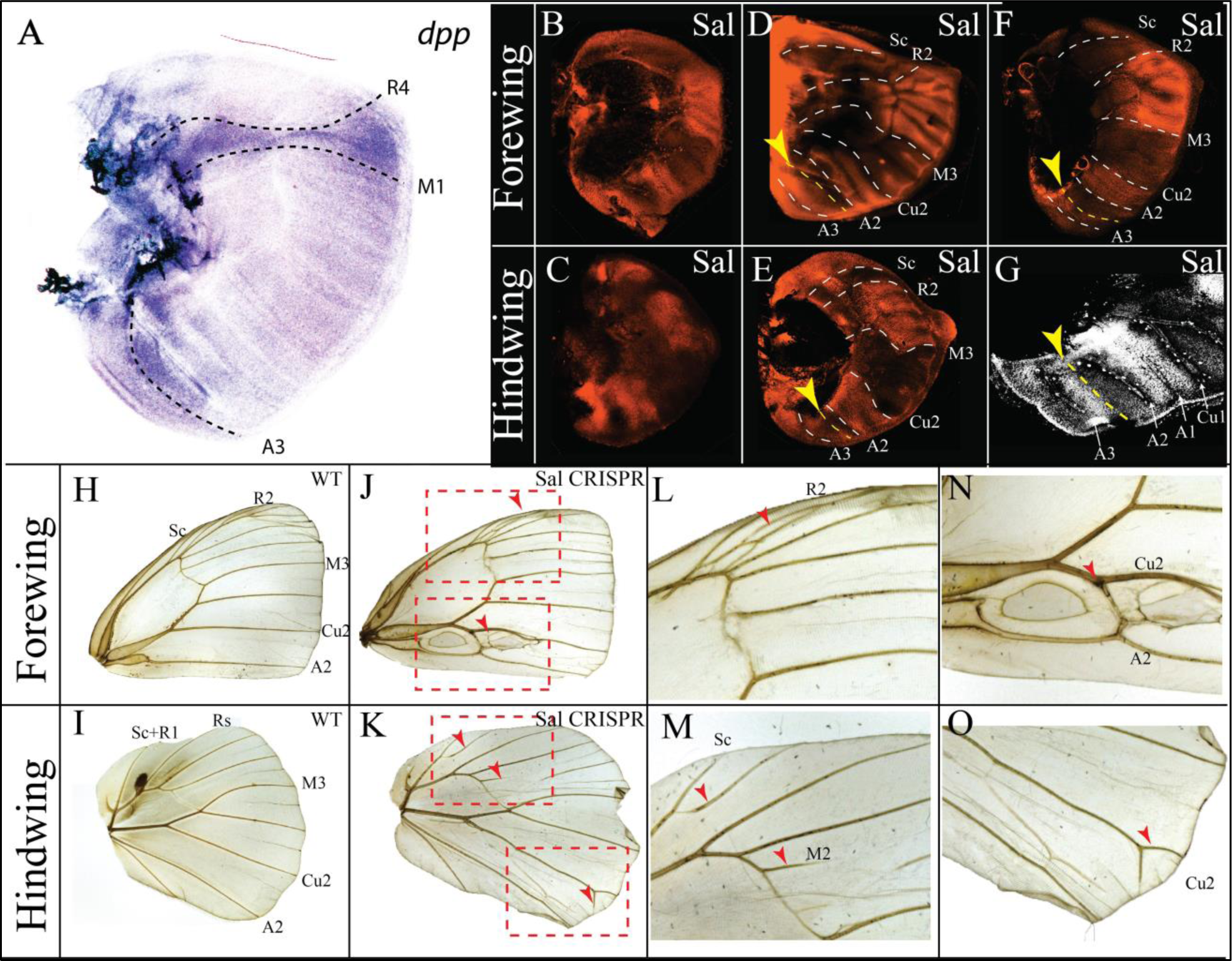
Expression of *dpp* in *Bicyclus anynana*, expression of Sal in *Bicyclus anynana* and *Pieris canidia*, and function of *sal* in *Bicyclus anynana*. **(A)** Expression of *dpp* in the larval wing of *Bicyclus* is visible at the A-P boundary spanning the M1 and R4 vein (for vein positioning at the same stage refer to **Fig. S7**) and along the A3 vein. Expression of Sal in **(B)** early, and **(D)** mid forewings, and in **(C)** early and **(E)** mid hindwings, showing distinct four distinct domains of expression: From the anterior margin till the Sc vein; from the R2 to M3 vein; from the Cu2 to A2 vein; and, from a boundary in between the A2 and A3 veins, and the posterior wing margin. **(F)** Expression of Sal in the larval forewing of *Pieris*; **(G)** Closeup of Sal expression in the lower posterior compartment showing the anterior boundary of the fourth Sal domain (yellow arrowhead and dotted yellow line). **(H)** WT adult forewing and **(I)** hindwing. **(J, L and N)** In CRISPR-Cas9 *sal* knock-out mosaic individuals’ ectopic veins are produced within the boundaries of Sal expression in forewings and **(K, M and O)** ectopic as well as missing veins are produced within the Sal expression domains in hindwings.

### Expression and function of *spalt*

To localize the transcription factor Sal we performed immunostainings in larval wings of *B. anynana* and *P. canidia* (*29*). Sal is expressed in four clearly separated domains in both early (Fig. 2, B and C) as well as in mid fifth instar wing discs (Fig. 2, D and E). The first domain appears anterior to the Sc vein. The second domain spans the interval between R2 and M3 veins. The third domain appears between the Cu2 and A2 veins. No expression is observed in a few cells posterior to the A2 vein and finally, a fourth domain appears posterior to a boundary between the A2 and A3 veins (Fig. 2G). These expression domains are also observed in *Pieris* (Fig. 2F).

To test the function of Sal in vein development we targeted this gene using CRISPR-Cas9. The phenotypes observed support a role for Sal in establishing vein boundaries at three of the four domains described above (Fig. 2, B-G). We observed both ectopic and missing vein phenotypes in both the forewing and the hindwing at the domains where Sal was present during the larval stage of wing development (Fig. 2, H-O). In the forewing, *sal* crispants generated ectopic and loss of vein phenotypes in between the R2 and the M3 vein domain (Fig. 2, J and L; Fig. S2. D and E) and ectopic veins between the Cu2 and A2 veins (Fig. 2N). In the hindwing, we observed ectopic veins connecting to the existing Sc vein (Fig. 2M), missing veins in between the Rs and M3 vein and ectopic veins connecting to the existing Cu2 vein (Fig. 2O).

### Expression of *bls*

To localize vein and intervein cells we performed *in-situ* hybridization of the intervein marker and vein suppressor gene *blistered* (*bls*). In the larval forewing and hindwing of *B. anynana, bls* is expressed in the intervein cells and lacks expression in the vein cells and cells around the wing margin (Fig. 3, A-C, Fig. S3, D-F).

**Fig. 3.**
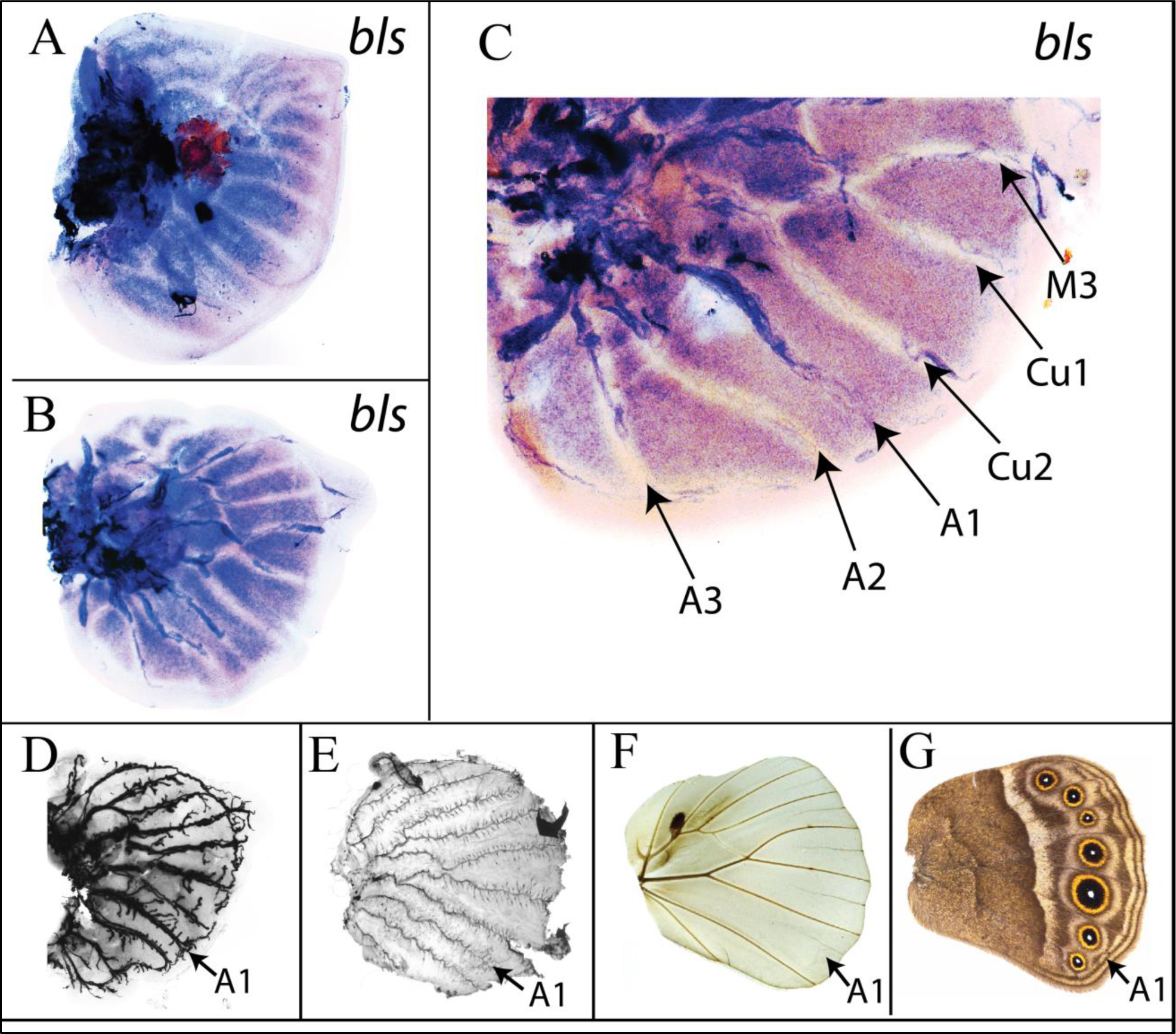
Expression of *blistered* (*bls*), and loss of the A1 vein in *Bicyclus anynana*. **(A)** Expression of *bls* in the larval forewing, and **(B)** larval hindwing of *Bicyclus*. *bls* is expressed in the intervein cells during larval wing development. **(C)** Closeup of *bls* expression in the posterior compartment where expression is present along the A1 vein. **(D-G)** Disappearance of the A1 vein during *Bicyclus* wing development. **(D)** Larval Wing; **(E)** Pupal wing; **(F)** Adult wing with scales removed; **(G)** Adult wing with scales. The A1 vein is observed in the larval and pupal stages, but it disappears in the adult stage.

## Discussion

A positional-information mechanism like that observed in *Drosophila* appears to be involved in positioning the veins in *Bicyclus* (and *Pieris*) but differences exist between flies and butterflies at multiple stages of the vein patterning mechanism (Fig. 4). These differences are highlighted below.

**Fig. 4.**
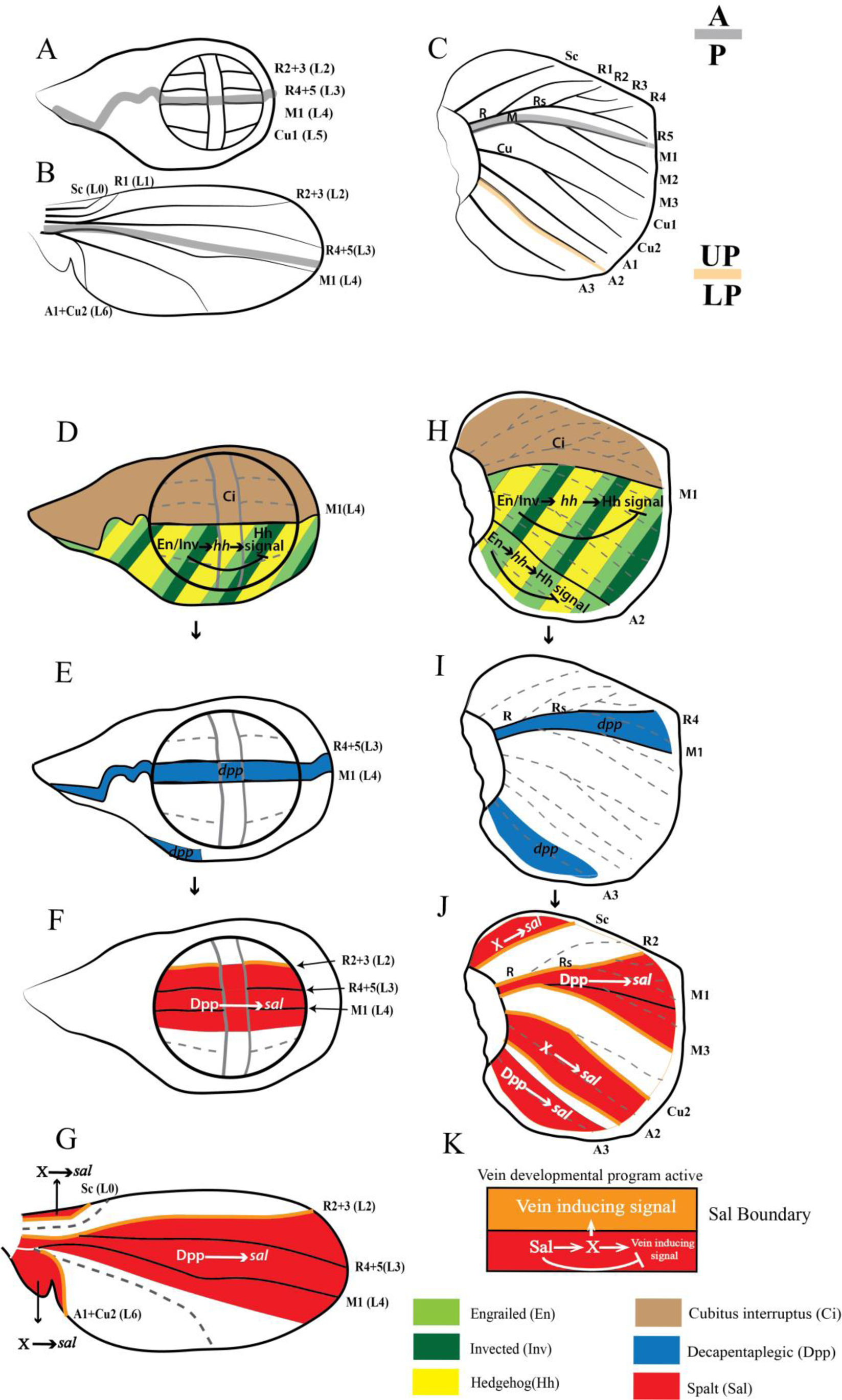
A model for venation patterning in *Bicyclus anynana* and comparison with *Drosophila melanogaster*. **(A)** Venation in *Drosophila* larval wing; **(B)** Venation in *Drosophila* pupal wing; **(C)** Venation in *Bicyclus* larval forewing; The Anterior-Posterior (A-P) boundary is marked by the thick grey line. The boundary between the Upper-Posterior (UP: from the M1 to A2 vein) and Lower-Posterior (LP: from the A2 vein till the posterior boundary) sectors of the wing is marked by the thick orange line. **(D-G)** Venation patterning in *Drosophila* wing (for details refer to the main text). **(H)** In *Bicyclus*, venation patterning is initiated by En and Inv expressed in the posterior compartment. En/Inv or En activates Hh in the posterior compartment, while suppressing Hh signaling. **(I)** A small amount of Hh diffuses into the anterior compartment where due to the presence of Hh signal transducer Ci, activates *dpp* in a thin stripe of cells in between the R4 and M1 veins. A second domain of *dpp* is activated around the A3 vein via a Hh independent mechanism. **(J)** In *Bicyclus*, Sal is expressed in four distinct domains. The first three domains are involved in induction of veins at their boundaries. **(K)** In *Drosophila* Sal induces vein development via the activation of a short-range hypothetical signal (X) and the inhibition of the signal from X. A small amount of X diffuses outside of the Sal expression domain, where it activates vein-inducing genes.

### The early wings of *Bicyclus* are subdivided into three gene expression domains instead of two as in *Drosophila*

One of the earlier steps in vein patterning in *Drosophila* is the separation of the wing blade into distinct compartments via the expression of En/Inv in the posterior compartment (*7*). *In-situ* hybridizations against the separate transcripts of *en* and *inv* in *Bicyclus*, showed that *en* is expressed across the whole posterior compartment (in the whole region posterior to the M1 vein), as in *Drosophila*, whereas *inv* is expressed only in the most anterior region of the posterior compartment, anterior to the A2 vein. This presumably leads to the higher En/Inv protein levels observed anteriorly, and lower protein levels posteriorly. While the *en in-situ* results are new, the *inv* expression is consistent with that observed in a previous study of *J. coenia* (*23*). The *inv* expression pattern in butterflies is, thus, distinct from that of *Drosophila* where *inv* is expressed homogeneously throughout the posterior compartment (*8*). These differences in expression of *en* and *inv* between *Drosophila* and *Bicyclus* essentially set up two main domains of gene expression in *Drosophila* wings but three in *Bicyclus* wings: An anterior domain with no *en* or *inv* expression, a middle domain with both *en* and *inv*, and a posterior domain with *en* but no *inv*.

### Two *dpp* signaling domains are established in *Bicyclus* whereas a single domain is present in the wing pouch of *Drosophila*

The next step in venation patterning in *Drosophila* is the establishment of the main *dpp* organizer along a stripe of cells, in the middle of the wing pouch (*30*). This group of *dpp*-expressing cells is established just above the *en/inv* expressing cells, at the A-P boundary, where the M1+2 (L4) vein will differentiate (*30*, *31*). In *Bicyclus* we also observe a group of cells expressing *dpp* at the A-P boundary above the M1 vein (Fig. 4I). This *dpp* expression in *Bicyclus* is likely driven by Hh diffusing from the posterior compartment to the anterior compartment where *Cubitus interruptus* (Ci), the signal transducer of Hh signaling, is present (*21*, *25*). In *Bicyclus*, however, there is a second group of *dpp*-expressing cells around the A3 vein (Fig. 4I). This second *dpp* domain in *Bicyclus* is probably activated via a Hh-independent mechanism, since no Ci or Patched (the receptor of Hh signaling) expression is observed in the posterior compartment around the A3 vein in butterflies (*21*, *25*). In *Drosophila*, there is also a group of *dpp*-expressing cells outside the wing pouch, which are activated via a Hh independent mechanism (*32*) (Fig. 4E). These two groups of cells could be homologous.

### Four domains of Sal expression are established in *Bicyclus*, whereas a single domain is present in *Drosophila* larval wings

The expression of *dpp* activates the next step in venation patterning in *Drosophila* which involves the activation of *sal* expression some distance away from the signaling center in a single main central domain (*6*, *9*, *11*, *33*). Here, the anterior boundary of Sal expression is involved in setting up the R2+3 (L2) vein (*6*, *10*, *11*, *34*). In *Bicyclus*, Sal is expressed in four separate domains in the larval wing, only two of which straddle the two *dpp* expression domains (Fig. 4, I and J). This suggests that Dpp might be activating *sal* in two of the domains where *dpp* and Sal are co-expressed and overlap, but a ligand other than Dpp, yet to be discovered, might activate *sal* in the first and third domains of Sal expression in *Bicyclus*. It is also interesting to note that in *Drosophila* only one Sal central domain is present during the larval stage, when vein differentiation is taking place, but a more anterior and a more posterior Sal expression domain appear during the pupal stage (*10*, *35*, *36*). To our knowledge no study has yet elucidated what drives the expression of Sal in these additional domains in *Drosophila* pupal wings.

### *sal* crispants show that three Sal boundaries are involved in positioning veins in *Bicyclus*, whereas a single Sal boundary performs this function in *Drosophila*

Sal knockout phenotypes in *Bicyclus* led to disruptions of veins in three out of the four Sal expression domains suggesting that, as in *Drosophila*, Sal is involved in setting up veins. *sal* crispants displayed: 1) ectopic Sc veins at the posterior boundary of the first Sal expression domain (Fig. 2M); 2) both ectopic and missing veins in the region of the second Sal domain straddling the A-P boundary, on both forewings and the hindwings, consistent with previous results on *Drosophila* (*10*, *37*); and 3) ectopic veins in both the forewing and the hindwing in the region of the third Sal domain (Fig. 2, J and K; Fig. S2, D and E). The final Sal expression domain in *Bicyclus* is present posterior to a boundary running in between the A2 and A3 vein (Fig. 2G), and we obtained no crispant with disruptions in veins in this area. Our data provide evidence, thus, that Sal boundaries of expression in domains 1, 2, and 3, are involved in differentiating veins at those boundaries in *Bicyclus*, whereas the boundary of the last Sal domain most likely are not used to position veins in the most posterior wing region.

The presence of both ectopic veins as well as disrupted veins in the domains of Sal expression in *Bicyclus* might be due to the disruption of the vein-intervein network in those regions. In *Drosophila*, ectopic and disrupted veins in *sal* knockout mutants are produced due to ectopic and missing *rho* expression (*10*). A proposed mechanism for how these genes interact involves Sal activating a hypothetical short-range diffusible protein (X) in the intervein cells and at the same time inhibiting the intervein cells from responding to the signal (*6*, *10*). A small amount of this diffusible protein moves towards the *sal-*negative cells activating vein inducing signals which includes genes such as *rho* (*10*) (Fig. 4K). Knockout of *sal* in clones of cells within a *sal-* expressing domain will create novel or missing boundaries of Sal+ against Sal-cells and will result in ectopic or missing expression of *rho*, activating or inhibiting vein development. In *Bicyclus*, we do not have direct evidence of *rho* expression, however, *bls* which has a complementary expression pattern to *rho* in *Drosophila* (*14*, *15*) is expressed throughout the *Bicyclus* larval wing with the exception of vein cells and the wing margin (Fig. 3, A-C), similarly to its expression pattern in *Drosophila*. This indicates that *rho* is likely expressed in the vein cells and that knocking out *sal* might result in ectopic or loss of *rho* in *Bicyclus* wing, resulting in ectopic and disrupted vein phenotypes (Fig. 2, J and K).

### Loss of *sal* expression domains and of the second *dpp* organizer likely led to venation simplification in *Drosophila*

Insect wing venation has simplified over the course of evolution, but it is unclear how exactly this simplification took place. Insect fossils from the Carboniferous period display many longitudinal veins in their wings compared to modern insects such as *Drosophila* or even *Bicyclus* (*1*, *38*, *39*). Many of the differences in venation remaining between *Bicyclus* and *Drosophila* are due to the additional loss of veins in the posterior compartment in *Drosophila* (Fig. 4, B and C). Sal expression domains and crispant phenotypes in *Bicyclus* indicate that the third Sal expression domain, present in *Bicyclus* but absent in *Drosophila* is involved in the formation and arrangement of posterior veins Cu2 and A2. In *Drosophila*, there is partial development of the Cu2+A1 (L6) vein and there are no A2 and A3 veins (Fig. 4B). The partial and missing veins in the posterior compartment of *Drosophila* are likely due to the reduction of the third and loss of the fourth Sal expression domains. The loss of the fourth Sal domain was perhaps a consequence of the partial loss of the second *dpp* organizer, and the reduction of the third domain mediated by the delayed expression of a yet undiscovered organizer in this region, that becomes activated only during the pupal stages in *Drosophila*.

### Simplification of venation is also achieved via silencing of vein inducing or vein maintenance mechanisms

Vein number reduction via loss of *sal/dpp* expression domains is one mechanism of vein reduction across evolution, but a different mechanism of vein reduction appears to take place downstream of the stable expression of these two genes. For instance, in *Bicyclus*, we observe the development of veins at both the boundaries of the second Sal domain (i.e., veins R2 and M3) (Fig. 4J), whereas in *Drosophila*, only cells abutting the anterior boundary of the homologous Sal expression domain activate the R2+3 (L2) vein (*10*) (Fig. 4F). Vein activation proceeds via the activation of vein-inducing genes such as *rho*, which does not take place at the posterior boundary of Sal expression in *Drosophila* (Fig. 4, F and G) (*10*). In *Bicyclus*, veins are also not being activated at the anterior boundary of the fourth Sal expression domain (just anterior to the A3 vein) (Fig. 4J). It is still unclear why veins don’t form at some boundaries of *sal* expression, but the paravein hypothesis proposes that loss of a vein inducing program at these boundaries, resulted in venation simplification in modern insects such as *Drosophila* (*6*).

Further venation simplification might be happening via disruptions of vein maintenance mechanisms, where vein induction is later followed by vein loss. In *Drosophila*, the maintenance of vein identity involves the stable expression of *rho* and the exclusion of *bls* from vein cells throughout wing development (*11*, *14*). Disruptions to this mechanism, however, appear to be taking place at the A1 vein during *Bicyclus* wing development (Fig. 3, D-G). The A1 vein is present during larval and early pupal wing development (Fig. 3, D and E) but is absent in adults (Fig. 3, F and G). In *Bicyclus*, *bls* is absent from the A1 vein in young larval wing discs (Fig. 3B, Fig. S3D). However, as the wing grows the expression of *bls* becomes stronger at the A1 vein (Fig. 3C; Fig. S3, E and F). It is possible that during late larval and pupal wing development *bls* becomes stably expressed at the A1 vein and the vein disappears as a result. It is unclear how the balance between *bls* and presumably *rho* expression is altered during development in the A1 veins of *Bicyclus*, but such a mechanism is contributing to the loss of that vein in adults and could be contributing to vein loss, in general, across insects.

In conclusion, we have provided evidence for the presence of three main domains of gene expression in the early wings of *Bicyclus*, an anterior, middle, and posterior domain instead of two (anterior and posterior) domains, as observed in *Drosophila*. We have discovered a second *dpp* expression domain in *B. anynana* present in the posterior of the wing disc, which is absent in *Drosophila* wing pouch where the vein patterning happens. Furthermore, we have described and functionally characterized four domains of Sal expression in *Bicyclus*, whose boundaries map to the development of multiple longitudinal veins in this species. Two of the Sal domains straddle the two *dpp* expression domains, and may be activated by a *dpp* gradient, but a different and yet undiscovered ligand (or ligands) is activating the two other Sal domains. The data presented in this study supports a Positional-Information mechanism involved in venation patterning in Lepidoptera as that observed in *Drosophila*. Moreover, the data provide support to the hypothesis of venation simplification in insects via loss of gene expression domains, silencing of vein inducing boundaries (*6*, *35*), and disruptions to vein maintenance programs (*11*). However, the mechanisms proposed in this paper cannot explain every feature of insect venation. Insects with left-right wing differences in their longitudinal vein branching patterns, such and in *Orosanga japaonicus* (*40*), and cross-vein patterns, such as in *Athalia rosae* (*41*, *42*) and *Erythremis simplicicolis* (*43*), most likely pattern their wings using both Positional-Information as well as Reaction-Diffusion mechanisms. Future comparative gene expression studies along with venation patterning modeling should continue to illuminate the evolution and diversity of venation patterning mechanisms in insects.

## Materials and Methods

### Animal husbandry

*Bicyclus anynana* butterflies were reared at 27°C in 12:12 day: night cycle. The larvae were fed young corn leaves and the adults mashed bananas.

### *In-situ* hybridization

Fifth instar larval wings were dissected in cold PBS and transferred into 1X PBST supplemented with 4% formaldehyde for 30 mins. After fixation, the wings were treated with 1.25 µl (20 mg/ml) proteinase-K (NEB, P8107S) in 1 ml 1X PBST and then with 2 mg/ml glycine in 1X PBST. Afterward, the wings were washed three times with 1X PBST, and the peripodial membrane was removed using fine forceps (Dumont, 11254-20) (in preparation for *in situ* hybridization). The wings were then gradually transferred into a pre-hybridization buffer (see Table S3 for composition) by increasing the concentration in 1X PBST and incubated in the pre-hybridization buffer for one hour at 60°C. The wings were then incubated in hybridization buffer (see Table S3 for composition) supplemented with 100ng/µl of probe at 60°C for 16-24 hrs. Subsequently, wings were washed five times with preheated pre-hybridization buffer at 60°C. The wings were then brought back to room temperature and transferred to 1X PBST by gradually increasing the concentration in the pre-hybridization buffer and they were later blocked in 1X PBST supplemented with 1% BSA for 1 hr. After blocking, wings were incubated in 1:3000 anti-digoxygenin labeled probe diluted in block buffer. To localize the regions of gene expression NBT/BCIP (Promega) in alkaline phosphatase buffer (See Table S3 for composition) was used. The wings were then washed, mounted in 60% glycerol, and imaged under a Leica DMS1000 microscope using LAS v.4.9 software.

### Immunostainings

Fifth instar larval wings were dissected in cold PBS and immediately transferred into a fixation buffer supplemented with 4% formaldehyde (see Table S4 for composition) for 30 mins. Afterward, the wings were washed with 1X PBS and blocked for one to two days in block buffer (see Table S4 for composition) at 4°C. Wings were incubated in primary antibodies against En/Inv (1:20, mouse 4F11, a gift from Nipam Patel, (*28*)), and Sal (1:20000, guinea-pig Sal GP66.1, (*29*)) at 4°C for one day, washed with wash buffer (see Table S4 for composition) and stained with secondary antibodies anti-mouse AF488 (Invitrogen, #A28175) and anti-Guinea pig AF555 (Invitrogen, #A-21435) at the concentration 1:500. The wings were then washed in wash buffer, mounted on an in-house mounting media (see Table S4 for composition), and imaged under an Olympus fv3000 confocal microscope, Zeiss Axio Imager M2.

### CRISPR-Cas9

Knock-out of *sal* was carried out using a protocol described previously(*44*). Briefly, a guide was designed against a 20 bp region targeting exon 1 of *sal* (see Supp file for sequence). The guide was tested via an *in-vitro* cleavage assay prior to injection. A total of 863 *Bicyclus* embryos were injected with 300 ng/µl of guide and 300 ng/µl of Cas9 protein (NEB; Cat. no.: M0641) mixed together in equal parts (total volume = 10 µl) with an added small amount of food dye (0.5 µl). The hatchlings were transferred into plastic cups and fed young corn leaves. After pupation, each individual was assigned a separate emergence compartment (a plastic cup with lid). Once eclosed, the adults were frozen at −20°C and imaged under a Leica DMS1000 microscope using LAS v4.9 software. Descaling of the adult wings for imaging was done using 100% Clorox solution (*45*). Mutant individuals were tested for insertions or deletions via an *in-vitro* endonuclease assay on the DNA isolated from the wings and then sequenced (Fig. S2).

## Acknowledgments

This work was supported by a Ministry of Education, Singapore grant MOE2015-T2-2-159 and the Department of Biological Sciences LHK fund GL 710221. TDB was supported by a Yale-NUS scholarship. We would like to thank Jocelyn Wee for rearing *Pieris canidia*, Kenneth McKenna and Fred Nijhout for initial discussions on the paper, Nipam Patel for the anti-Engrailed/Invected antibody and, Jocelyn Wee, Suriya Narayanan Murugesan and Sofia Sigal-Passeck for useful comments. The authors declare no conflict of interest.

## Supplementary Materials

**Fig. S1.**
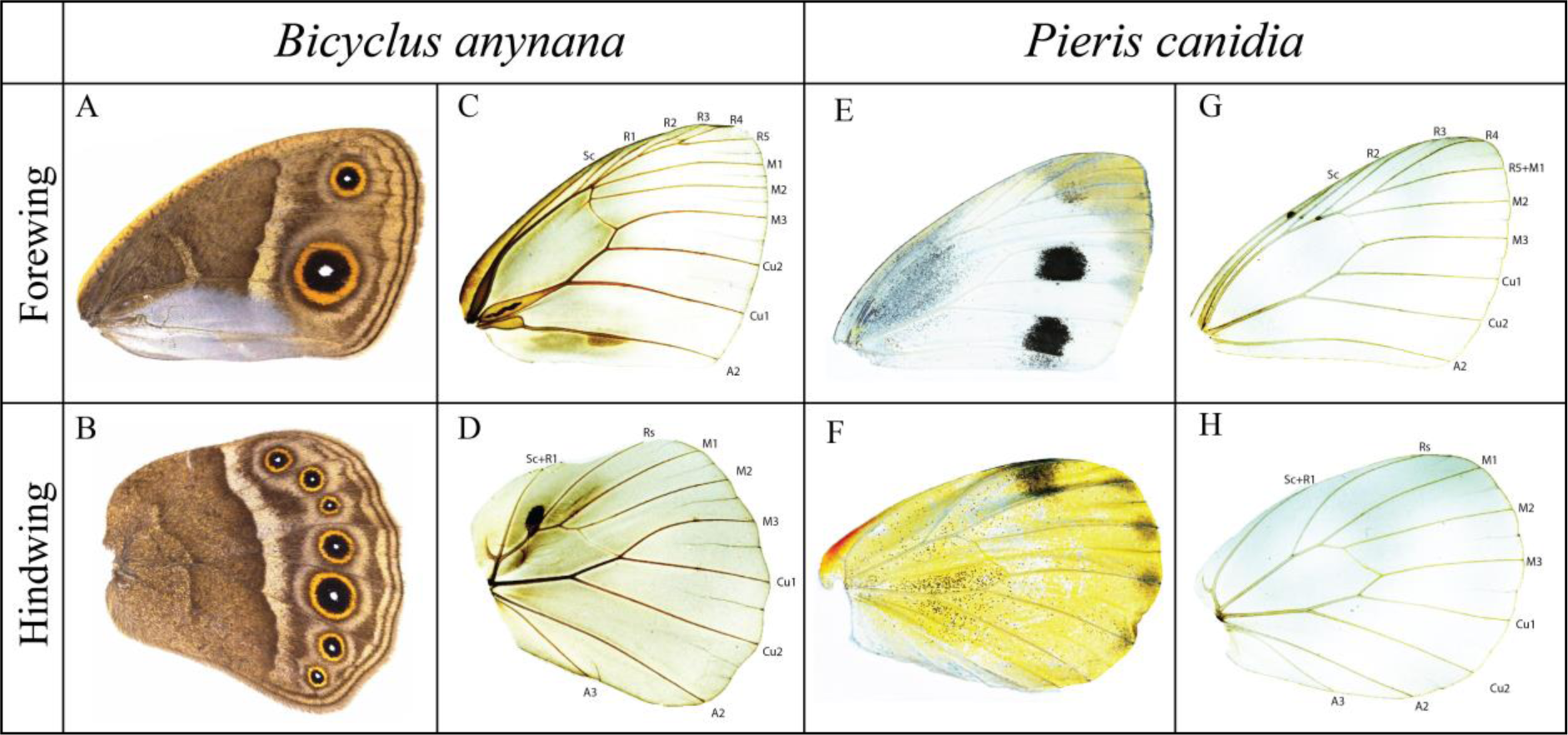
Venation pattern in butterflies. **(A and C)** *Bicyclus anynana* forewing, **(B and D)** and hindwing. **(E and G)** *Pieris canidia* forewing, **(F and H)** and hindwing.

**Fig. S2.**
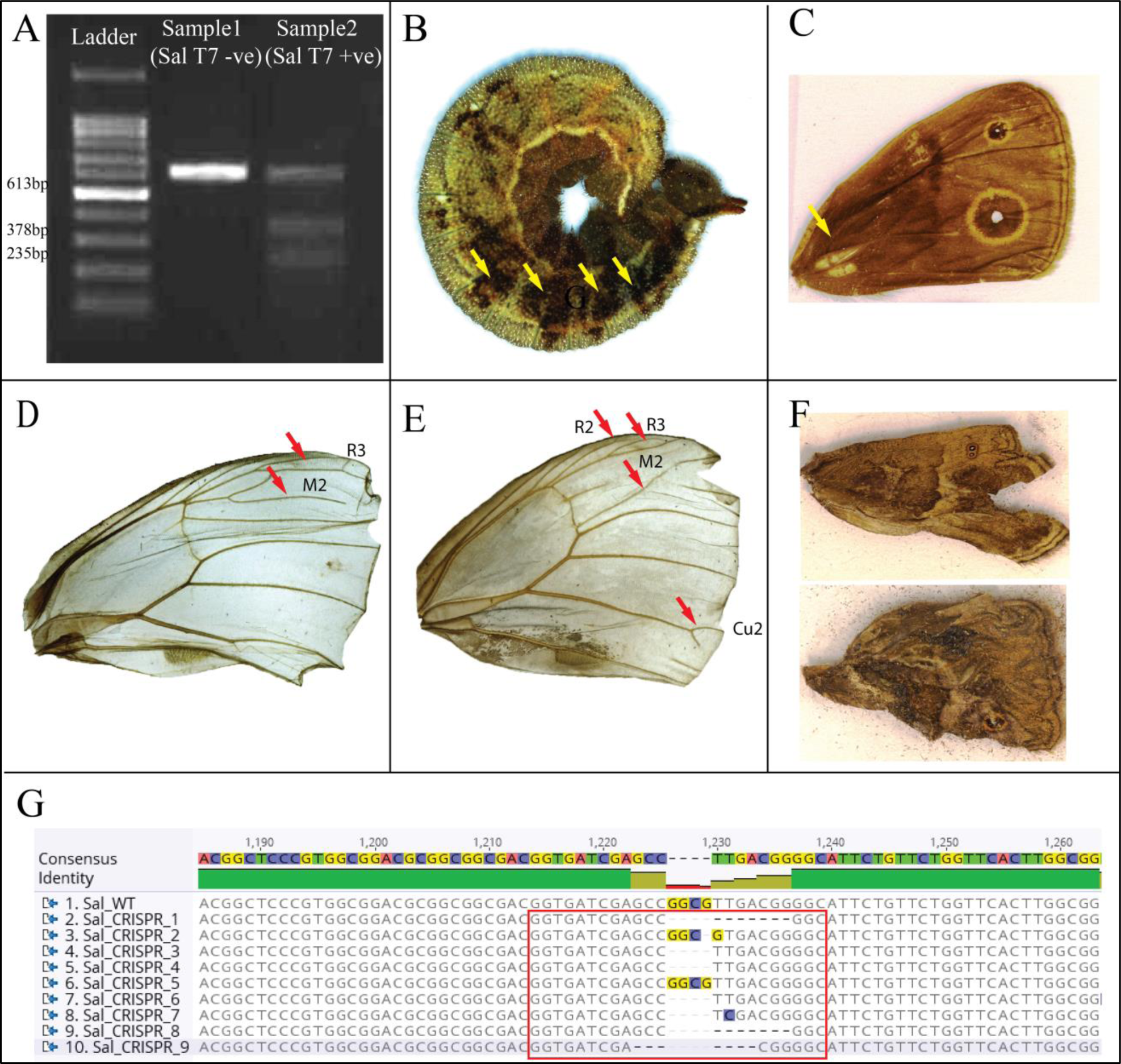
Spalt CRISPR-Cas9 on *Bicyclus anynana* butterflies. (**A**) T7 endonuclease assay on *sal* guide and Cas9 injected individuals. Sample 2 with T7 endonuclease added shows two shorter DNA bands indicating cleavage of the PCR product. (**B**) Pigmentation defects on the embryos. (**C**) Adult wing showing defects in the auditory organ and eyespot on the wing, (**D and E)** Descaled adult forewings showing venation defects in the region where Sal is expressed (red arrows). (**F**) Severe wing patterning defects in some individuals were observed. (**G**) Deletions of nucleotides at the CRISPR-Cas9 target site. The target region is marked by the red box.

**Fig. S3.**
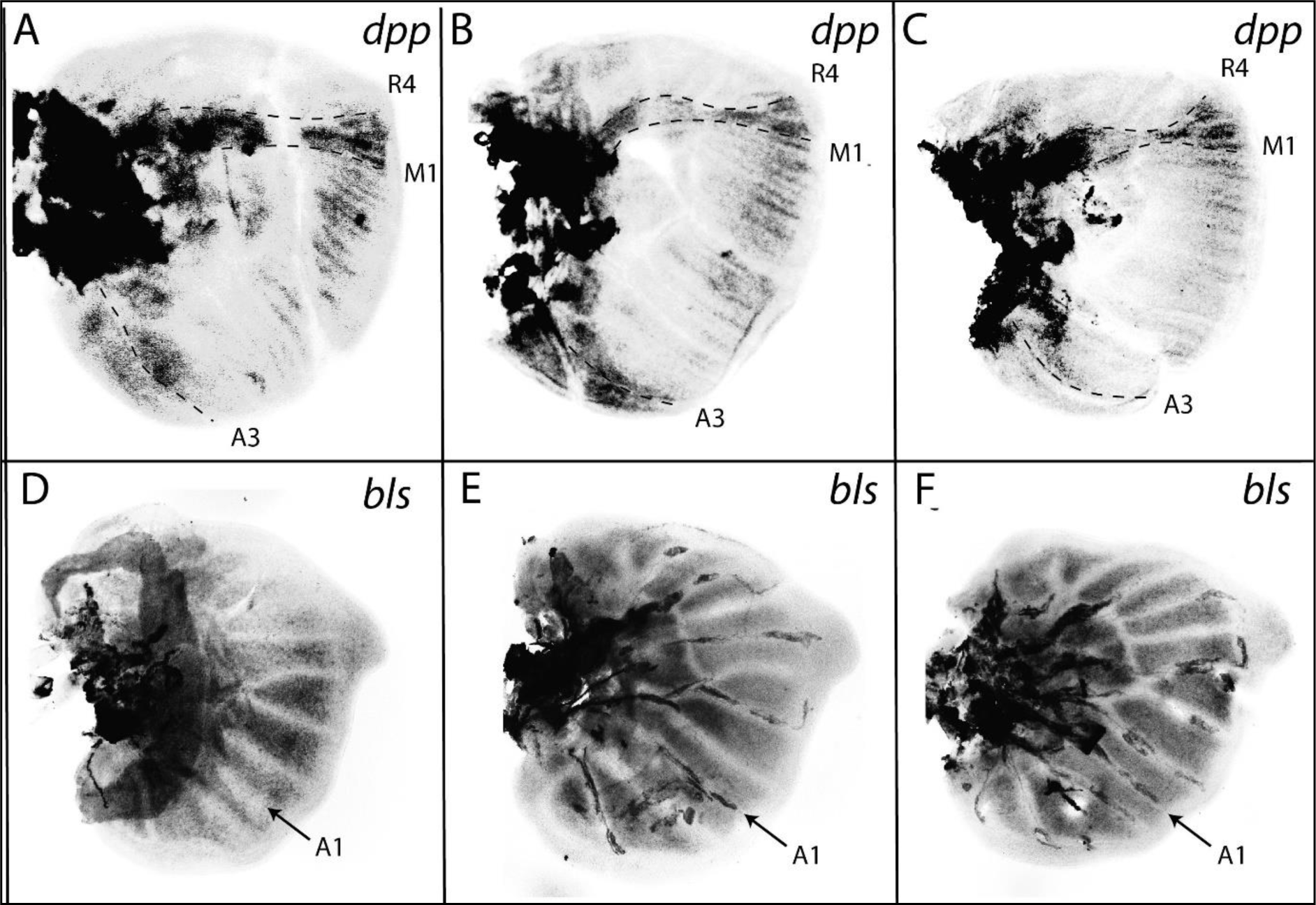
*decapentaplegic* (*dpp*) and *blistered* (*bls*) mRNA localization in *Bicyclus anynana*. (**A-C**) *dpp* is expressed in two domains in the larval wings. (**D-F**) Expression of *bls*. *bls* is absent at the A1 vein at an early stage. However, during later stages *bls* has a stronger expression at the A1 vein.

**Fig. S4.**
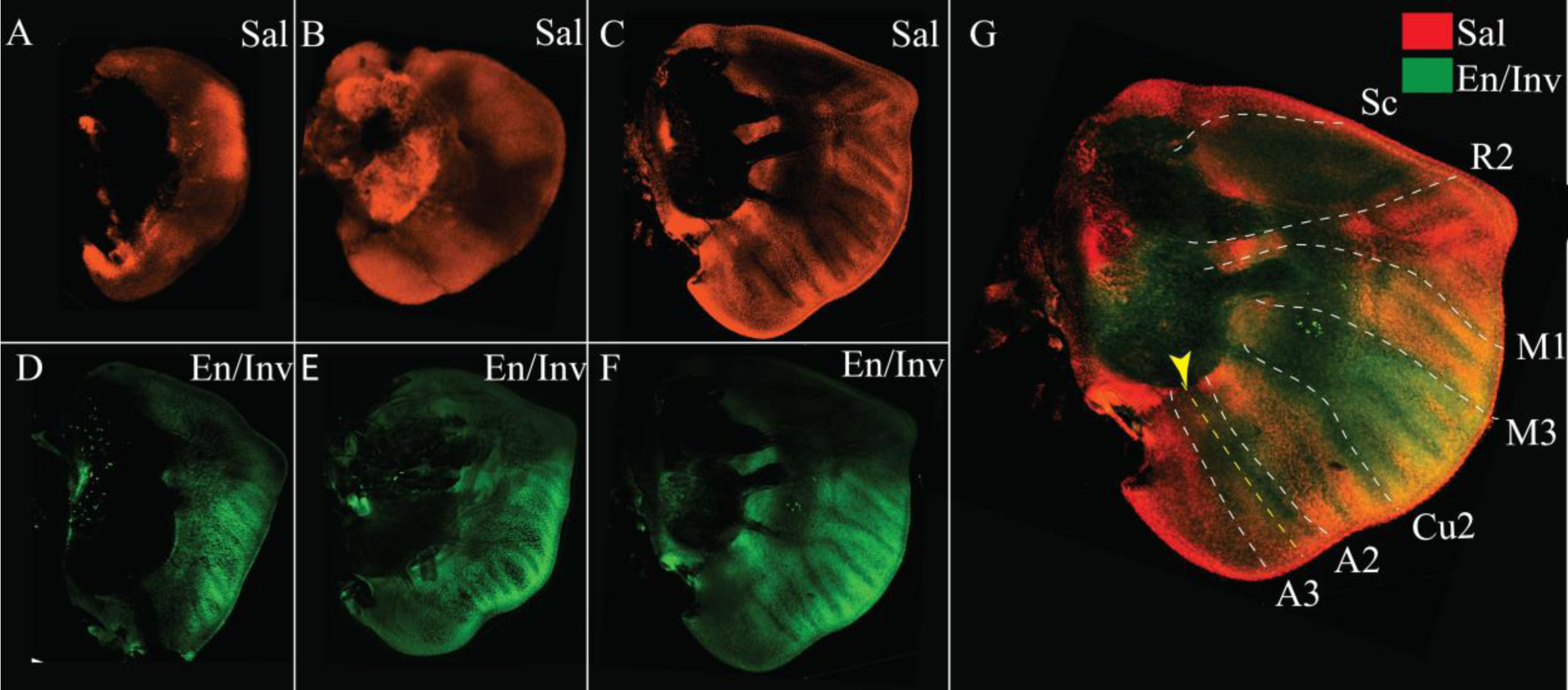
Sal and En/Inv in *Bicyclus anynana*. (**A-C**). Sal staining at different stages of wing growth. (**D-F**). En/Inv staining at different stages of wing growth. (**G**). Co-staining of Sal and En/Inv.

**Fig. S5.**
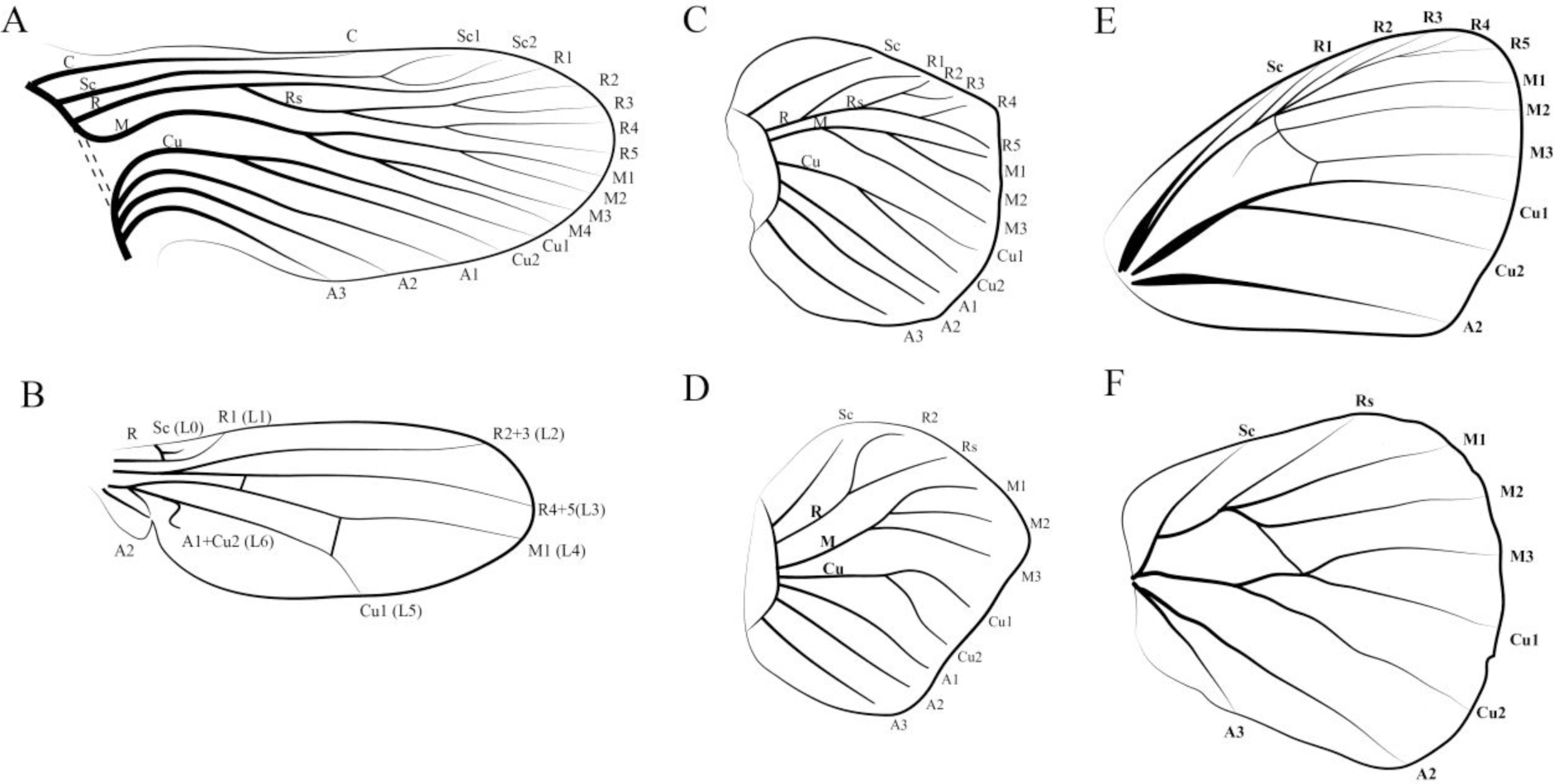
Venation patterns in insects. **(A)** Comstock-Needham hypothetic venation of primitive insects (redrawn from (*2*)), **(B)** Wing venation of *Drosophila melanogaster* (redrawn from (*11*)), **(C)** Larval forewing venation and **(D)** hindwing venation of *Bicyclus* butterflies. Larval wings of *Bicyclus* were drawn based on methylene blue staining’s (Fig. S5). **(E)** Adult forewing and **(F)** hindwing venation of *Bicyclus*.

**Fig. S6.**
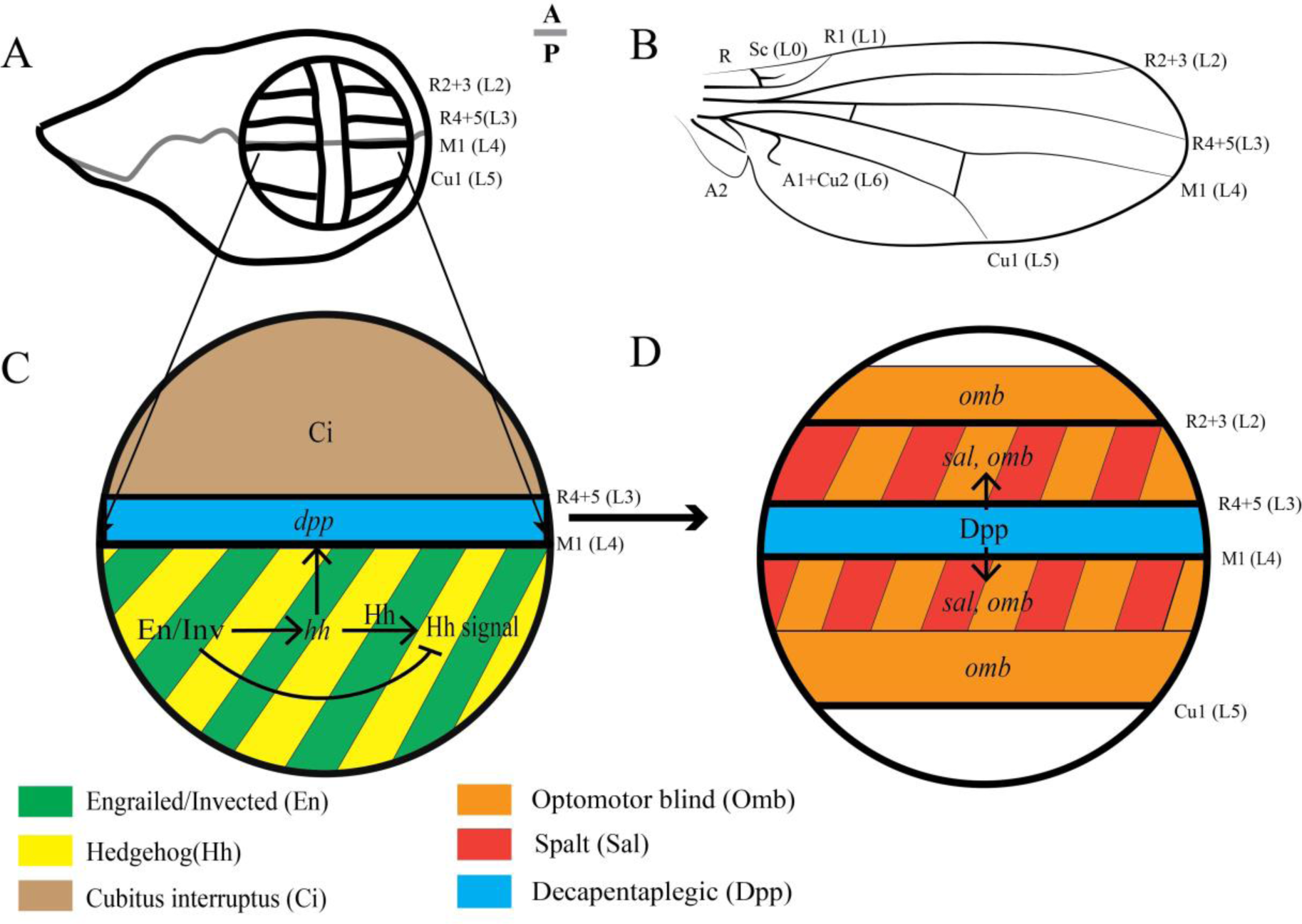
Molecular mechanism involved in venation patterning in *Drosophila melanogaster*. **(A)** Larval wing disc of *Drosophila*. During the larval stage, the wing is divided into two populations of immiscible cells belonging to the Anterior (A) and Posterior (P) compartments. The boundary where these two-populations meets is referred to as the Anterior-Posterior (A-P) boundary (marked by the gray line). **(B)** Adult wing of *Drosophila*. **(C)** Venation patterning is initiated by the transcription factors En and Inv in the posterior compartment that activate expression of *hh* while suppresing Hh signaling. Hh is a short-range diffusible morphogen. A small amount of Hh diffuses into the anterior compartment where the presence of Ci activates the BMP ligand *dpp*. The veins R4+5 (L3) and M1 (L4) form at the anterior and posterior boundary of the Hh signaling domain where *dpp* is expressed. **(D)** Dpp protein then acts as a long-range morphogen activating both *spalt* (*sal*) and *optomotor-blind* (*omb*) at high concentrations, and only *omb* when the concentration falls below the *sal*-inducing threshold. The vein R2+3 (L2) develops at the anterior boundary of Sal, while the vein Cu1 (L5) develops at the posterior boundary of Omb protein domains.

**Fig. S7.**
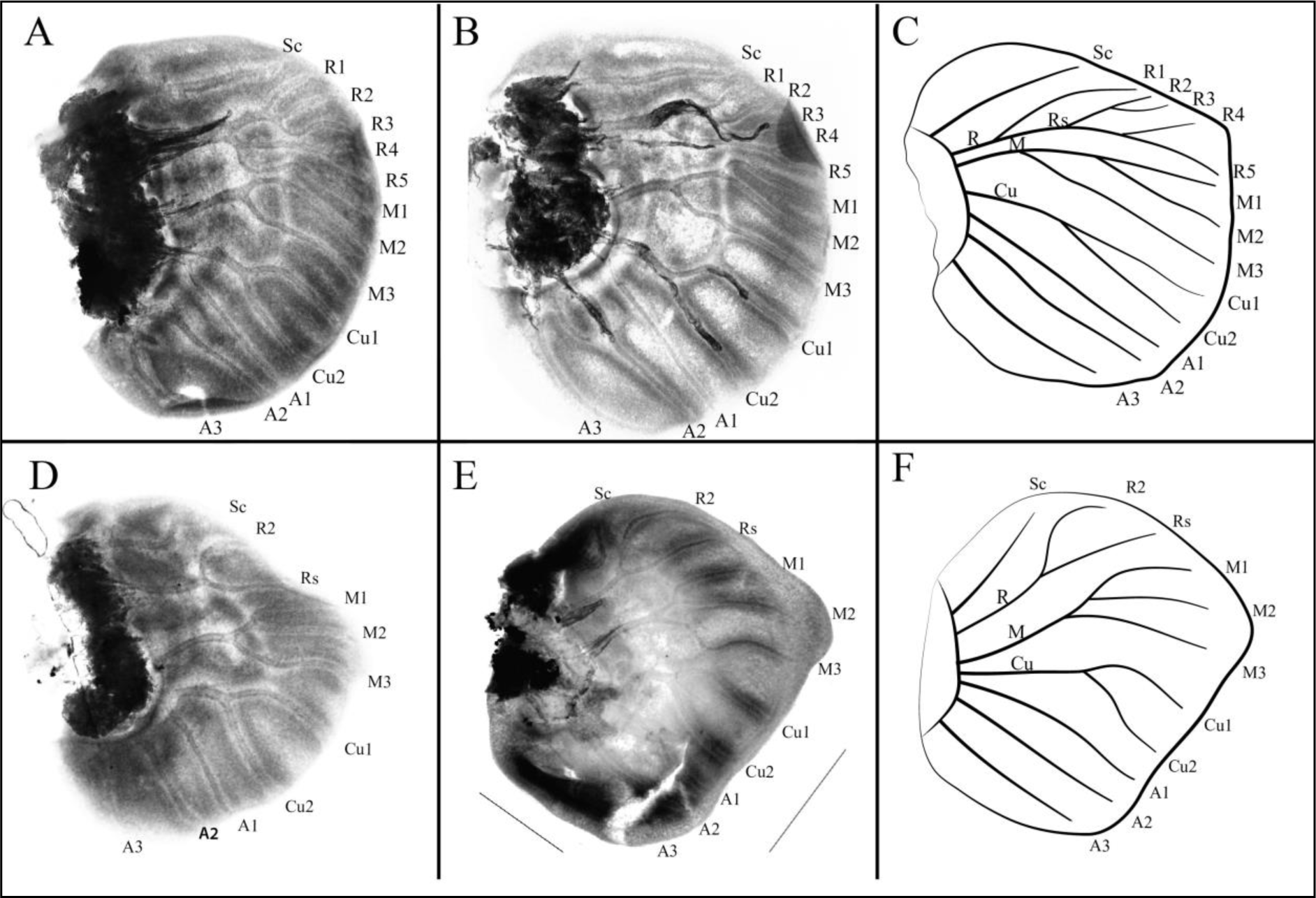
Methylene blue staining of *Bicyclus anynana* larval wings. **(A, B)** Forewing stained with methylene blue; **(D, E)** Hindwing stained with methylene blue; **(C)** Illustration of forewing venation; **(F)** Illustration of hindwing venation.

**Table S1.** Spalt CRISPR-Cas9 injection table.

| Sl. No. | Concentration | Date | Eggs Injected | Hatchlings | % Hatchlings |
| --- | --- | --- | --- | --- | --- |
| 1. | 300 ng/μl | 28th Sept 2018 | 302 | 48 | 15.9 |
| 2. | 300 ng/μl | 10th Oct 2018 | 306 | 25 | 8.2 |
| 3. | 300 ng/μl | 11th Nov 2018 | 120 | 18 | 15 |
| 4. | 300 ng/μl | 9th Feb 2019 | 135 | 8 | 5.9 |

**Table S2.**
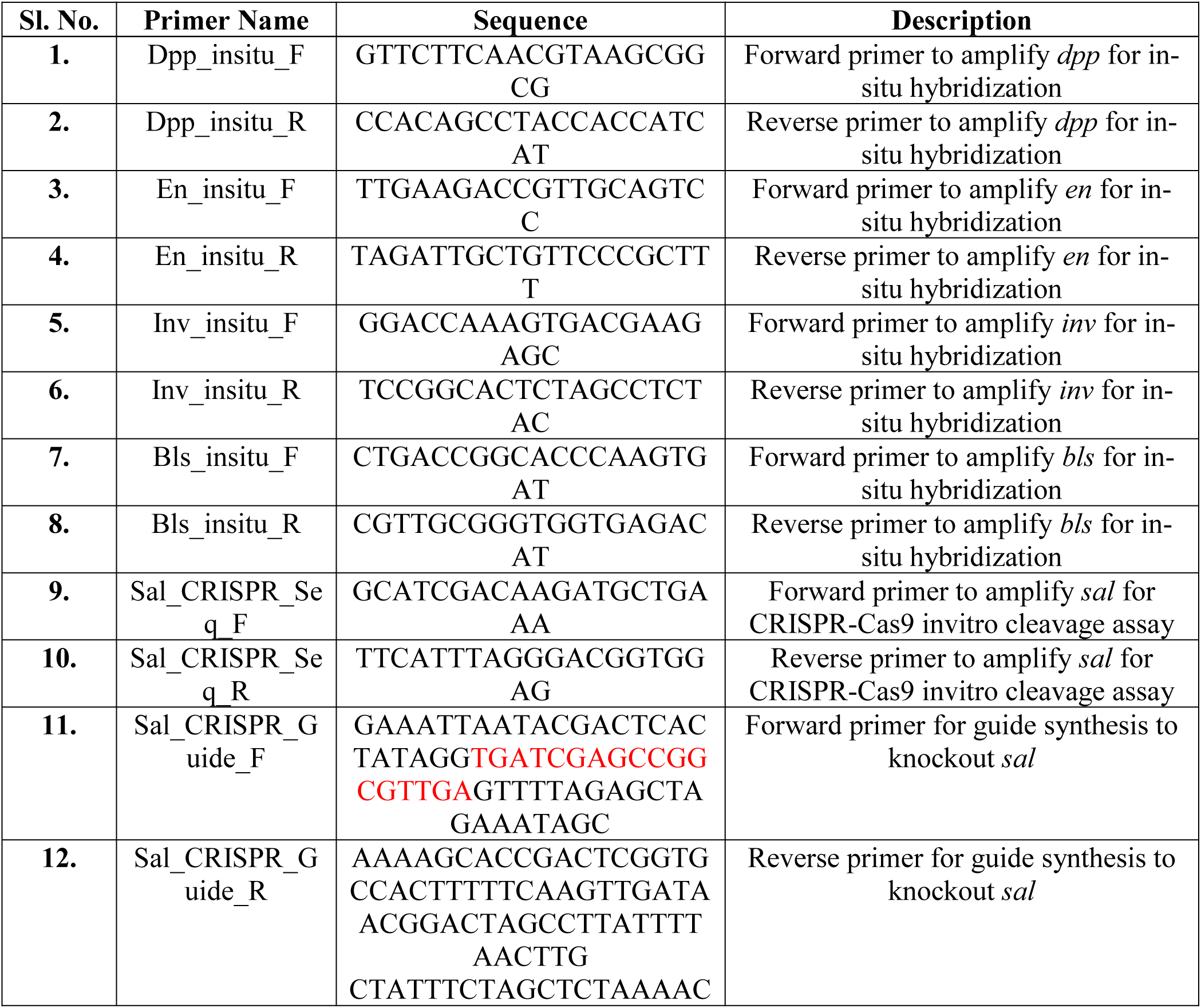
Primer table.

**Table S3.**
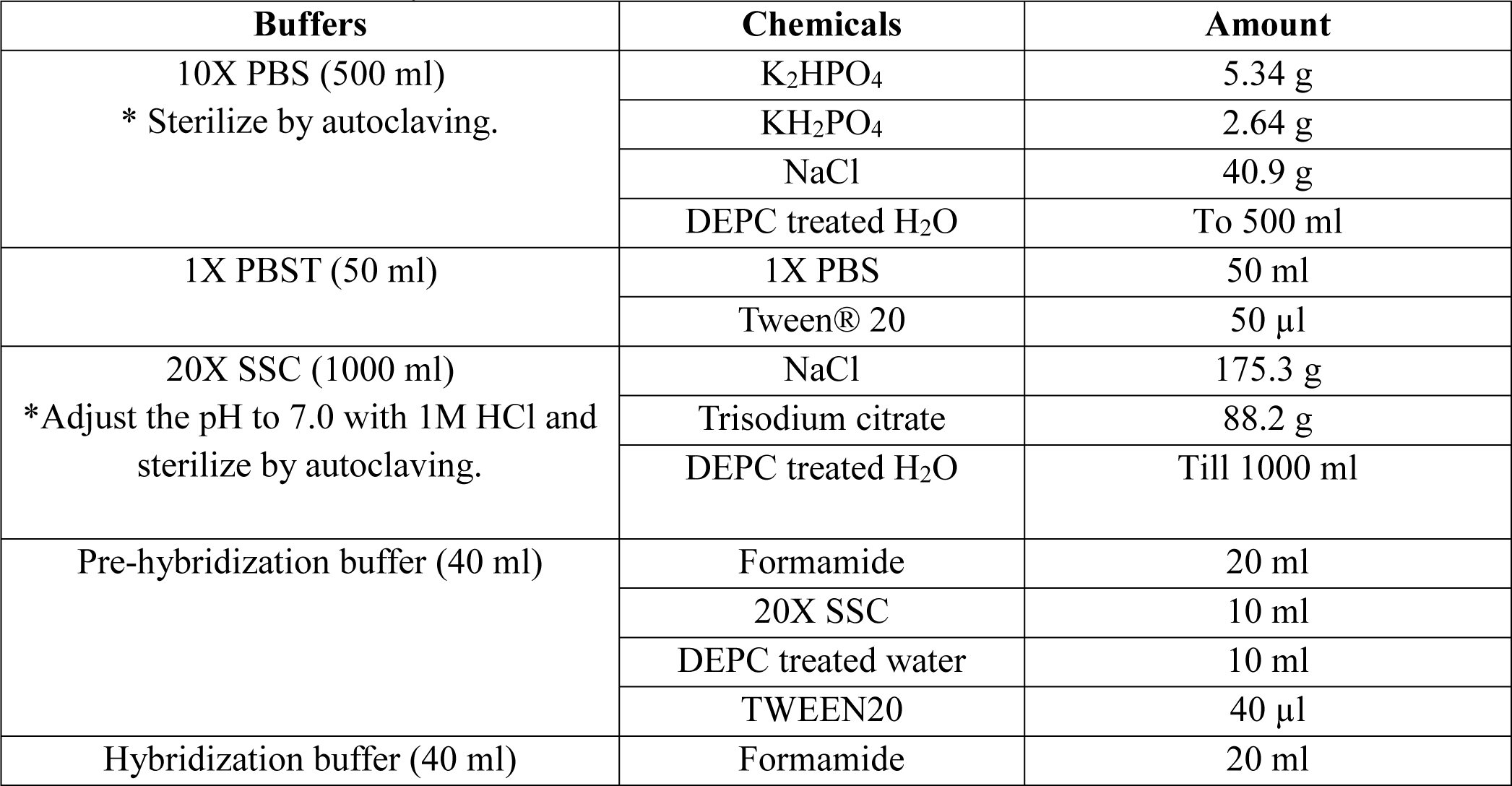
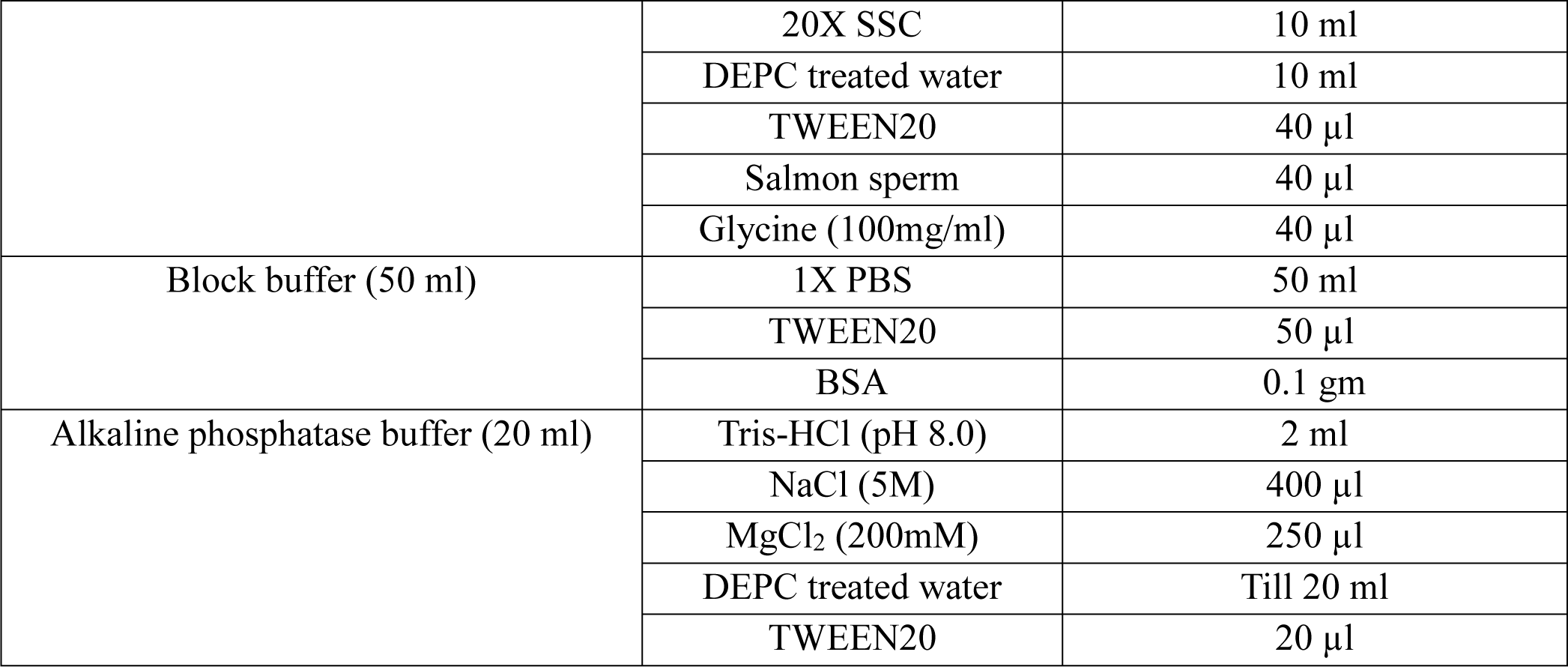
*In-situ* hybridization Buffers.

| Buffers | Chemicals | Amount |
| --- | --- | --- |
| 10X PBS (500 ml)<br>* Sterilize by autoclaving. | K<sub>2</sub>HPO<sub>4</sub> | 5.34 g |
|  | KH<sub>2</sub>PO<sub>4</sub> | 2.64 g |
|  | NaCl | 40.9 g |
|  | DEPC treated H<sub>2</sub>O | To 500 ml |
| 1X PBST (50 ml) | 1X PBS | 50 ml |
|  | Tween® 20 | 50 µl |
| 20X SSC (1000 ml)<br>*Adjust the pH to 7.0 with 1M HCl and sterilize by autoclaving. | NaCl | 175.3 g |
|  | Trisodium citrate | 88.2 g |
|  | DEPC treated H<sub>2</sub>O | Till 1000 ml |
| Pre-hybridization buffer (40 ml) | Formamide | 20 ml |
|  | 20X SSC | 10 ml |
|  | DEPC treated water | 10 ml |
|  | TWEEN20 | 40 µl |
| Hybridization buffer (40 ml) | Formamide | 20 ml |
|  | 20X SSC | 10 ml |
|  | DEPC treated water | 10 ml |
|  | TWEEN20 | 40 µl |
|  | Salmon sperm | 40 µl |
|  | Glycine (100mg/ml) | 40 µl |
| Block buffer (50 ml) | 1X PBS | 50 ml |
|  | TWEEN20 | 50 µl |
|  | BSA | 0.1 gm |
| Alkaline phosphatase buffer (20 ml) | Tris-HCl (pH 8.0) | 2 ml |
|  | NaCl (5M) | 400 µl |
|  | MgCl<sub>2</sub> (200mM) | 250 µl |
|  | DEPC treated water | Till 20 ml |
|  | TWEEN20 | 20 µl |

**Table S4.** Immunohistochemistry Buffers.

| Buffers | Chemicals | Amount |
| --- | --- | --- |
| Fix buffer (30 ml) | M PIPES pH 6.9 (500 mM) | 6 ml |
|  | mM EGTA pH 6.9 (500mM) | 60 µl |
|  | % Triton x-100 (20 %) | 1.5 ml |
|  | mM MgSO<sub>4</sub> (1M) | 60 µl |
|  | 37% Formaldehyde | 55 µl per 500 µl of buffer |
|  | dH<sub>2</sub>O | 22.4 ml |
| Block buffer (40 ml) | 50 mM Tris pH 6.8 (1 M) | 2 ml |
|  | 150 mM NaCl (5 M) | 1.2 ml |
|  | 0.5% IGEPAL (NP40) (20%) | 1 ml |
|  | 5 mg/ml BSA | 0.2 gr |
|  | H<sub>2</sub>O | 35.8 ml |
| Wash buffer (200 ml) | 50mM Tris pH 6.8 (1 M) | 10 ml |
|  | 150 mM NaCl (5 M) | 6 ml |
|  | 0.5% IGEPAL (20 %) | 5 ml |
|  | 1 mg/ml BSA | 0.2 gr |
|  | dH<sub>2</sub>O | 179 ml |
| Mounting media | Tris-HCl (pH 8.0) | 20 mM |
|  | N-propyl gallate | 0.5% |
|  | Glycerol | 60% |

### Sequence of *engrailed* used for *in-situ* hybridization

TTGAAGACCGTTGCAGTCCGAACCAGGCCAACAGCCCCGGTCCGGTCACCGGCAGAG TCCCTGCGCCTCACTCCGAAGTAAGAAACGNGTACCAAAGTCAATACACTTGCACGA CTATCGATCAAAGGTTTGACAGAACGATGACAGTGGTGAAAGTGCAGCCGAATTCAC CACCGATGAGTCCACTGACGTGAAGCCCATAATCCCTGAGTTTGAAGACAAGAGAAA CCGACAACCACCACCAACCATACCCTTCTCTATCAGCAACATATTACACCCAGAATTC GGTTTGACAGCGATTCGAAAAACGAACAAAATCGAAGGACCAAAACACGTCGGCCC CAACCACAGCATTTTGTACAAACCTTATTTGTCGAACGAGTTATCGAGTTCGAAATTC AATTTCGATTATTTAAAATCTAAGGATGATTTCGGTGCATTACCTCCACTTGGCGGTT TGAGGCAGACCGTGTCGAATATTGGAGAACAGAAGGAGGCACCAAAGATTATAGAG CAGCAGAAGAGGCCAGATTCAGCCAGCTCTATTGTCTCTTCCACATCTAGCGGGGCT TTATCGACGTGTGGCAGCACTGACGCCAACAGCAGTCAAAGCGGGAACAGCAATCTA

### Sequence of *invected* used for *in-situ* hybridization

GGACCAAAGTGACGAAGAGCACGACCCCTACTCGCCCAACACTAGAGACACCATCACACCAGACTTCATAGAAGAAGACAAACAAGACAGGCCTATACACACATCCTCTTTCTCCATACACAATGTCCTTAAGAAGGAAAGAGACAGTAATAGTCCTGAGAACGTCTTCTCAACTGAAAAGTTGTTGCAAAGTACACCGAACTTTGAAGATTCTAGGAACTCTGAAAGCGTTAGTCCGAGACTTGAAGATGATCACAATGAAAGAGCTGATATAAGTGTTGATGACAACTCTTGTTGTAGTGATGATACTGTGCTATCTGTTGGCAATGAAGCCTTACCAACCAATTACCCAAACGACAAAGATCCGAACCAAGGCTTAACCTCCTTCAAACATATACAAACTCATTTGAACGCAATATCACAGTTAAGTCAAAATTTAAACATAAACCAACCAATCCTCCTACGACCCAACCCAATAACACCAAACCCGTTAATGTTCCTAAACCAACCGTTGTTATTCCAAAACCCTTTAATAAACCAAGTGGATTTAAAATCAGGGTTACCGAGAATCGGCTTGCAGCAAAACAATTTAAATTTGAACCAAAATTACATGAATTATGCGAGAAAAAATGAACTGAACGAAAGACGACAGAGTTATTCACCGAAGTTACATGAAAATGAGTCAAGTAGAGATTTTATTAACCAAGGATGTTTGAAATTTAGCATTGATAATATACTGAAAGCTGATTTTGGTAGACGAATTACTGATCCGTTGACAAAGAGAAAAACGAAGACGAGGCAGTATGAGGCAAAATCTACCCCTGTCAAAGAGGTTCAGTCTCCCCCTAAAGAGGTAGAGGCTAGAGTGCCGGA

### Sequence of *decapentaplegic* used for *in-situ* hybridization

GTTCTTCAACGTAAGCGGCGTACCGGCCGACGAGGTGGCGCGCGGCGCCGACCTCTCGTTCCAACGAGCCGTCGGCACCACCGGCAGACAGAGACTGTTGTTGTACGACGTGGTGCGCCCTGGCCGCCGCGGCCACTCCGAGCCGATCCTGCGGCTGCTGGACTCCGTTCCGCTCCGGCCCGGGGAGGGAATCGTCAACGCCGACGCTCTGGGAGCGGCGCGACGGTGGCTCAAAGAGCCCAAACATAATCACGGACTATTAGTGCGAGTGTTAGAAGAAGACGCCGCGAGTGCGAGCAGGGACGCGAAGTTCCCGCACGTGCGCGTGCGCAGACGCGTCACGGACGAGGAGGAGGAGTGGCGGACGGCGCAGCCGCTGCTCATGCTGTACACGGAGGACGAGCGCGCGCGCGCGTCGCGGGAGACGAGCGAGCGGCTGACGCGCAGCAAGCGCGCGGCGCAGCGGCGGGGGCACCGCGCGCACCACCGCCGCAAGGAGGCGCGCGAGATCTGCCAGCGCCGCCCGCTGTTCGTCGACTTCGCGGACGTGGGCTGGAGCGACTGGATCGTGGCCCCGCACGGCTACGACGCGTACTACTGCCAGGGCGACTGCCCCTTCCCGCTGCCGGACCACCTCAACGGCACGAACCACGCGATAGTGCAGACTCTGGTCAACTCAGTGAACCCCGCGACGGTGCCCAAAGCGTGCTGCGTGCCGACGCAACTCTCATCTATATCTATGTTATATATGGACGAAGTGAACAATGTGGTGCTTAAAAACTATCAGGACATGATGGTGGTAGGCTGTGG

### Sequence of *blistered* used for *in-situ* hybridization

GCATACGAGCTATCAACGCTGACCGGCACCCAAGTGATGCTGCTGGTCGCGTCGGAGACCGGCCACGTGTACACGTTCGCGACACGCAAACTGCAGCCGATGATCACGTCCGACTCGGGCAAGCGGCTCATACAGACGTGCCTCAACTCGCCCGACCCGCCCACCACCAGCGAGCAGCGCATGGCCGCCACCGGCTTCGAGGAGACCGAGCTCACGTATAACGTTGTAGACGACGAGATGAAGGTGAGACAACTGGCGTACGCTAACCAGTACCCCATAGAGCACCACCCGGGGTTGGCGCCGTCGCCACTGCAGCAGTACCACCAGCACCCGCCCTGCCCCTCGCCCCTCCCCCTCGGCTCGCTGGGCCAGCCGTACTCGCACGCGCATCTATCGCACCCCCACATGTCTCACCACCCGCAACG

### Region of *spalt* used for CRISPR-Cas9 (Highlighted in red)

GCATCGACAAGATGCTGAAAATAATAATAGTCTCGAAGACGGCGAGGCCGAAATACCTGAAGCCGACATGCCCCCCGTGGGTCTGCCGTTCCCTTTGGCAGGACACGTTACTCTTGAGGCTCTACAAAATACGAGAGTAGCGGTCGCCCAATTCGCTGCAACAGCGATGGCAAATAATGCGAATAACGAAGCTGCTATACAAGAATTACAAGTGTTACACAACACTCTATACACTTTACAGTCACAACAAGTATTTCAACTTCAGTTAATACGTCAGCTTCAGAATCAGTTATCTCTAACTCGACGGAAAGAAGACGATCCACACAGCCCACCGCCAAGTGAACCAGAACAGAATGCCCCGTCAACGCCGGCTCGATCACCGTCGCCGCCGCGTCCGCCACGGGAGCCGTCGCCTGTTATACCCTCTCCTCCTACTAGCCAAAGTTTGCCGTCGACTCACACACATCACACACCCAAAACTGAACAGATATCTATCCCTAAGATTCCAACTTCCTCACCATCTTTAATGACCCACCCACTTTATAGTTCAATTTCTTCGTCATTAGCATCTTCC ATCATAACAAACAATGATCCTCCACCGTCCCTAAATGAA

